# Altered age-dependent somatic mutational trajectories in ALS motor cortex neurons

**DOI:** 10.64898/2026.11.17.767052

**Authors:** Liyueyue Liu, Charlotte van Dijk, Sam Bonsall, Rodrigo Siqueira Kazu, Sai Zhang, Jonathan Cooper-Knock, Jan Veldink, Kevin Kenna

## Abstract

Amyotrophic lateral sclerosis (ALS) is a fatal neurodegenerative disease in which disease onset is influenced by genetic, age-associated and environmental factors. Somatic mutations accumulate throughout life and may provide a molecular link between aging and neuronal vulnerability. Here, we investigated neuronal somatic mutational patterns in ALS and control motor cortex using three previously generated 10X single-nucleus RNA and ATAC sequencing datasets. Within the genomic regions accessible to 10X sequencing, variant calls showed established features of neuronal somatic mutagenesis, including neuronal enrichment, age-associated accumulation and recovery of the clock-like SBS5 mutational signature. ALS was associated with a disease-specific age-dependent trajectory of neuronal somatic mutation burden, with higher burden in younger ALS donors and a decreasing difference with age. SBS5 represented the largest fitted mutational component and showed a corresponding age-dependent difference. A similar pattern was observed in an independent single-cell whole-genome sequencing dataset. These findings support age-dependent alterations in neuronal somatic mutational patterns in ALS.

## Introduction

Amyotrophic lateral sclerosis (ALS) is a fatal late-onset neurodegenerative disease characterized by progressive degeneration of motor neurons in the spinal cord and motor cortex, with an estimated lifetime risk of approximately 1 in 350 and a 5-year survival rate of 20–30% [1]. Although inherited genetic factors contribute substantially to ALS susceptibility, as supported by twin [2] and genome-wide association studies[3], 90%-95% ALS cases are sporadic and disease onset typically occurs in later adulthood [1]. This contrast between genetic contribution, sporadic presentation and age-dependent onset suggests that ALS may arise from interactions between inherited susceptibility, aging-related cellular processes and environmental exposures [4][5]. Somatic mutations accumulate throughout life in post-mitotic neurons[6] and provide a molecular record of endogenous and environmental mutagenic processes[7], making them a potential link between aging, genome maintenance and late-onset neurodegenerative disease[8].

In the healthy aging human brain, single-cell genome sequencing studies have shown that neurons acquire somatic mutations at an approximately linear rate with age [9,10,11,12,13].The clock-like mutational signature SBS5 represents a prominent component of this age-associated accumulation and has been repeatedly observed across brain regions and sequencing platforms [8,10,11,14,15]. Together with the other major clock-like signature, SBS1, SBS5 is widely observed in post-mitotic neurons [8,10,11,14,15]. However, unlike SBS1, the underlying etiology of SBS5 remains incompletely understood and is likely complex. SBS5 mutational patterns have been associated with various exogenous exposures and endogenous such as aging, tobacco smoking, nucleotide excision repair (NER) deficiency, and potentially multiple DNA damage and repair processes[16–18]. Neurons are particularly susceptible to the accumulation of somatic mutations because they are long-lived, post-mitotic cells and therefore depend largely on replication-independent DNA repair pathways, including nucleotide excision repair, base excision repair and non-homologous end joining [19]. In addition, their high metabolic activity and substantial oxygen consumption during neuronal signaling expose them to elevated levels of reactive oxygen species and associated oxidative stress [20]. Together, these features make somatic mutations a cumulative molecular record of the endogenous [21][22] and environmental processes [23] acting on the neuronal genome over time.

Recent high-resolution single-cell DNA sequencing studies have begun to map somatic mutation landscapes in the healthy aging brain and in neurodegenerative diseases, including Alzheimer’s disease [23][24], chronic traumatic encephalopathy [25] and ALS [26]. These approaches, including primary template-directed amplification (PTA)[9], NanoSeq [11], Hairpin Duplex Enhanced Fidelity sequencing [13] and other single-cell whole-genome sequencing methods [27–29], provide high-confidence mutation calls at single-neuron resolution. However, they remain limited by the number of donors and cells that can be profiled [10], making it difficult to resolve cohort-level disease-associated mutation trajectories across aging in neurons. Conversely, large-scale public single-cell 10X snRNA & snATAC datasets provide access to hundreds of thousands of single cells across many donors. Although sparse coverage restricts mutation detection to a limited genomic window, previous work has shown that neuronal somatic mutations are enriched in open chromatin and transcriptionally active regions [8][10][30][31], supporting the use of 10X-detectable regions as an informative window into broader neuronal somatic mutational trends. The integration of these two modalities provides complementary advantages, with snRNA-seq enabling robust cell-type annotation and snATAC-seq extending genomic coverage beyond the 3′-end regions captured by snRNA-seq while also avoiding RNA associated technical artefacts relating to RNA editing [32][33], variable transcript abundance [34], and post-mortem interval (PMI)-associated RNA degradation [35][36].

Here, we asked whether public 10X snRNA & snATAC datasets could be used as a cohort-scale framework to study neuronal somatic mutational aging in ALS. Across three ALS and control 10X single-nucleus motor cortex cohorts, we evaluated modality-specific covariates, benchmarked somatic variant detection within the effective snATAC-seq mutation-detectable genomic window, used snRNA-seq to annotate disease-associated neuronal states. We observed that 10X-derived calls recapitulated established features of neuronal somatic mutagenesis, including higher detectable mutation burden in neurons compared with oligodendrocytes and astrocytes, age-associated burden increases in healthy neurons and recovery of the clock-like mutational signatures SBS5 and SBS1. We further found that ALS was associated with a non-linear reshaping of the neuronal somatic mutational aging trajectory in the motor cortex, with SBS5 emerging as the major underlying mutational component. These findings position large-scale single-cell snRNA & snATAC data as a complementary approach for investigating disease-associated somatic mutational trajectories in the aging human brain. They further suggest that somatic mutation burden in ALS follows a more complex age-dependent trajectory than a simple linear increase with age, warranting genome-wide, high-resolution investigation of the somatic mutational landscape in larger cohorts with greater numbers of donors and cells.

## Results

### 1. snATAC captures global somatic mutation trends in neurons

We harmonized three previously generated 10X single-nucleus datasets from human motor cortex [37–39], comprising 118 ALS and control samples and 885,929 cells in total (Table 1). The Li and Bonsall cohorts included snATAC-seq and snRNA-seq data, whereas the Pineda cohort comprised snRNA-seq data. Raw reads were uniformly processed and aligned using Cell Ranger, followed by somatic variant calling with SComatic (Fig. 1a; Methods) [33]. Due to sparse coverage within fine-grained neuronal subtypes, neuronal subtypes were aggregated into a single neuronal compartment. Before testing for ALS-associated differences, we evaluated the validity of detected somatic mutations based on three established features of neuronal somatic mutagenesis: higher mutation burden in neurons relative to other brain cell types, continual accumulation with age and characteristic mutational signatures. Given the distinct genomic regions interrogated by snATAC-seq and snRNA-seq, as well as potential RNA-specific sources of variation, the two modalities were considered separately.

**Fig. 1.**
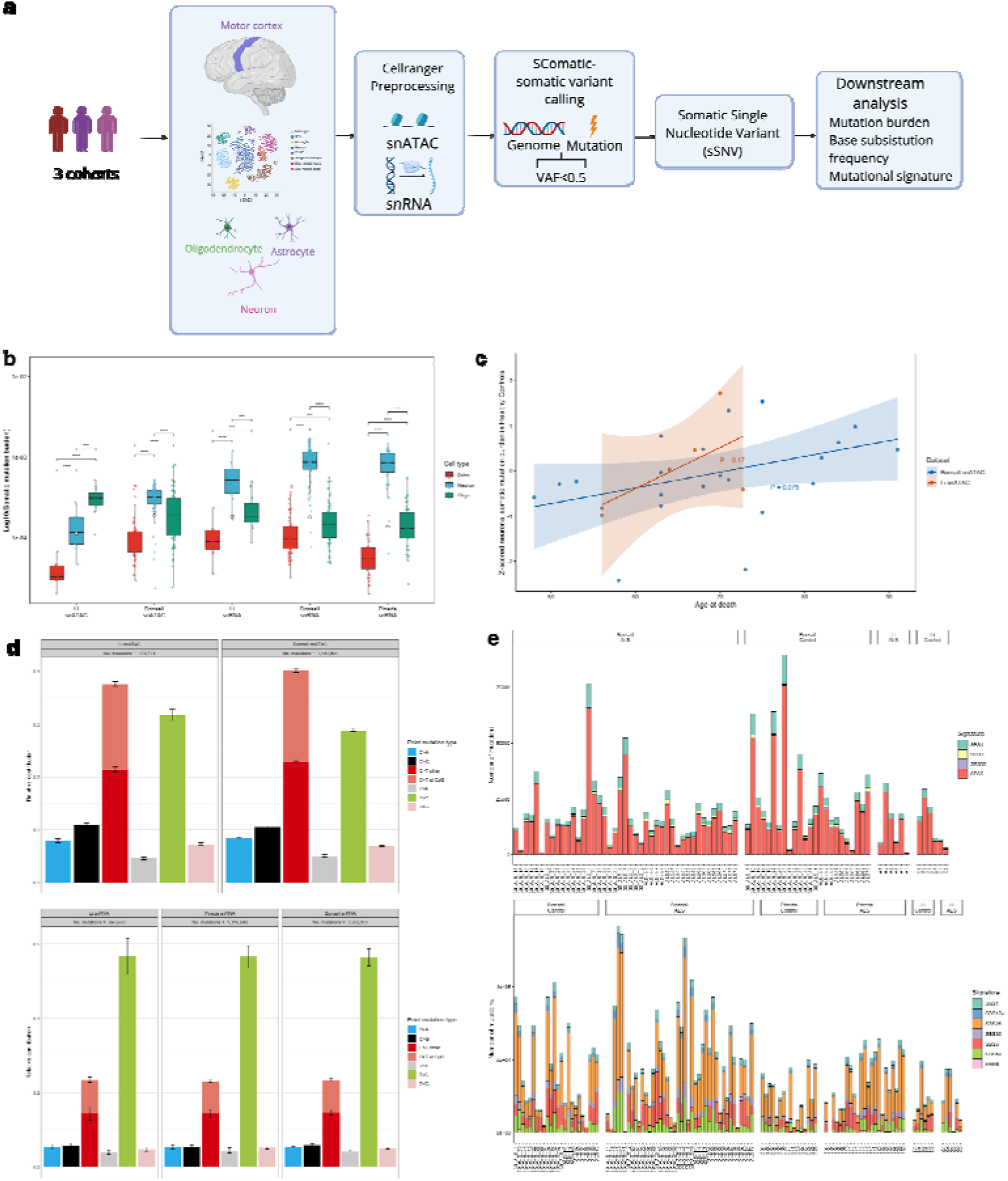
Study design and benchmarking of somatic variant detection from 10x single-nucleus sequencing data. **a**, Overview of the analytical workflow for processing, somatic variant calling, filtering and downstream analysis of motor cortex snATAC-seq and snRNA-seq data from three ALS cohorts, comprising 72 individuals with ALS and 46 controls (118 individuals; 885,929 cells). Data were preprocessed using Cell Ranger, and somatic single-nucleotide variants (sSNVs) were called using SComatic. Variants with a variant allele fraction ≥0.5 were excluded before analyses of somatic mutation burden, base-substitution frequencies and mutational signatures. **b,** Somatic mutation burden across astrocytes, neurons and oligodendrocytes in 2 snATAC-seq and 3 snRNA-seq datasets. Pairwise comparisons between cell types were performed separately within each dataset using two-sided Wilcoxon rank-sum tests with Benjamini–Hochberg correction. All pairwise comparisons remained significant after multiple-testing correction (adjusted P = 1.86 × 10-20 to 8.58 × 10 ). Boxes indicate the median and interquartile range; whiskers extend to 1.5 times the interquartile range, and points represent individual samples. Mutation burden is displayed on a log scale. **c,** Association between age at death and z-scored neuronal somatic mutation burden in healthy control samples from the Bonsall and Li snATAC-seq datasets. Linear regression lines and 95% confidence intervals are shown. A positive age-related trend was observed in both datasets, although neither association reached statistical significance (Bonsall, P = 0.075; Li, P = 0.17). **d,** Relative contributions of the six single-base substitution classes across the Li and Bonsall snATAC-seq datasets and the Li, Pineda and Bonsall snRNA-seq datasets. Bars represent the mean relative contribution of each substitution class, and error bars indicate 95% confidence intervals.e, COSMIC mutational signature exposures obtained by refitting the signatures retained through stepwise selection. Stacked bars show the estimated number of mutations attributed to each signature per sample, grouped by dataset and disease status, for snATAC-seq datasets (top) and snRNA-seq datasets (bottom).

**Table 1.** Sample information of different 10X cohorts used in this study.

| **Cohort** | **Sample number** | **Phenotypes** | **Type of disease** | **Age range** | **Sex** | **Cell number** | **Modalities** |
|---|---|---|---|---|---|---|---|
| Pineda | 40 | ALS:23<br>Control:17 | Sporadic:17<br>C9:6 | 52-79 | F:12<br>M:11 | 242,215 | snRNA |
| Li | 12 | ALS:6<br>Control:6 | C9:6 | 58-72 | F:3<br>M:3 | 95,675 | snRNA&snATAC |
| Bonsall | 67 | ALS:43<br>Control: 24 | Sporadic:25<br>C9:18 | 46-86 | F:29<br>M:38 | 548,039 | snRNA&snATAC |

First, we evaluated somatic mutations detected from the two snATAC-seq datasets. The Li dataset was notably smaller and had substantially fewer callable neuronal sites per donor than Bonsall (Extended Data Fig. 1a), reducing the number of mutations available for analysis. Nonetheless, neurons showed significantly(adjusted P = 1.86 × 10-20 to 8.58 × 10 , Supplementary Table 1) higher detectable mutation burdens than astrocytes in both datasets and higher burden than oligodendrocytes in the Bonsall dataset (Fig. 1b). In healthy controls, neuronal mutation burden increased with age in both datasets, although neither association individually reached statistical significance (Bonsall, P = 0.075; Li, P = 0.17; Fig. 1c). Base-substitution profiles were reproducible between the two snATAC-seq datasets, and mutational signature refitting using forward step selection [31][40] within selected COSMIC panels (Method, Extended Data Fig. 2a) [12,24,40–45] with MutationalPatterns[46] recovered the clock-like SBS5 signature previously associated with age-related somatic mutagenesis[47] in human neurons (Fig. 1d, e) [8,10,11,14,15] and other noncancerous tissues [48]. Thus, across cell-type distribution, age association and mutational signature profile, snATAC-derived variants showed patterns consistent with established features of neuronal somatic mutagenesis.

We next assessed the sensitivity of our method for somatic variant detection within the genomic regions accessible to snATAC-seq. SomatoSim [46–49] was used to introduce variants across a range of variant allele fractions (VAFs) [50] into neuronal snATAC-seq data, followed by the same SComatic calling workflow used for the observed data (Methods). Most covered sites had relatively low read depth (Extended Data Fig. 1b), resulting in strong VAF- and depth-dependent detection probabilities (Extended Data Fig. 1c, d). Among successfully simulated variants, 51.48% occurred at sites reaching ≥5 reads, the default minimum depth threshold for a callable site in SComatic. Together, these analyses thus indicate that snATAC-seq provides an incomplete but biologically informative survey of somatic mutations within its accessible genomic window.

Next, we evaluated variants detected by snRNA-seq. Neurons showed significantly higher detectable mutation burden than astrocytes and oligodendrocytes across the snRNA-seq datasets (Fig. 1b). However, base-substitution profiles differed substantially from those observed in snATAC-seq, with a pronounced enrichment of T>C (A>G) substitutions that was reproducible across snRNA-seq datasets (Fig. 1d). Although our analyses filtered out known RNA-editing sites [51–53], such an enrichment could potentially be explained by residual contributions from unannotated RNA editing sites or other RNA-specific processes. Alternatively, region-specific contributions from somatic mutations could also be relevant [54]. In either case, mutational signature refitting using the same method (Method, Extended Data Fig. 2b) applied to snATAC-seq data did not recover an SBS5-dominated pattern (Fig. 1e). We also observed no clear association or consistent direction of effect between snRNA-derived neuronal mutation burden and donor age across control cohorts (Extended Data Fig. 3a). Notably, PMI differed significantly between control cohorts showing opposite age-associated trends (P = 0.0269; Extended Data Fig. 3b). Consistent with a potential PMI-related effect, snRNA-derived neuronal mutation burden was significantly associated with PMI in the Pineda control samples after adjustment for donor age (P = 0.0346; Extended Data Fig. 3c). Given the apparent impact of RNA-specific sources of variation, subsequent analyses of somatic mutational trajectories were therefore restricted to snATAC-seq.

### 2. Altered age-dependent somatic mutational trajectories in ALS motor cortex neurons

We next asked whether ALS associates with changes in the age-associated accumulation of neuronal somatic mutations. In snATAC neurons, the relationship between mutation burden and age differed significantly between ALS and controls (Interaction P = 0.024) (Fig. 2a). The fitted trajectories suggested a non-linear pattern: ALS samples showed higher neuronal mutation burdens than controls before ∼64 years of age, whereas this difference diminished and reversed in older samples. The age-dependent pattern was concordant with recent single-cell PTA data from the ALS motor cortex [26], in which ALS neurons showed significantly higher somatic SNV burden among younger donors, whereas this difference was no longer apparent among older donors (Fig. 2b). This independent observation supports the biological plausibility of the changing ALS–control relationship across the age trajectory. These findings suggest that ALS is associated not merely with a uniform increase in neuronal somatic mutation burden, but with a reshaping of the age-associated mutational trajectory.

**Fig. 2.**
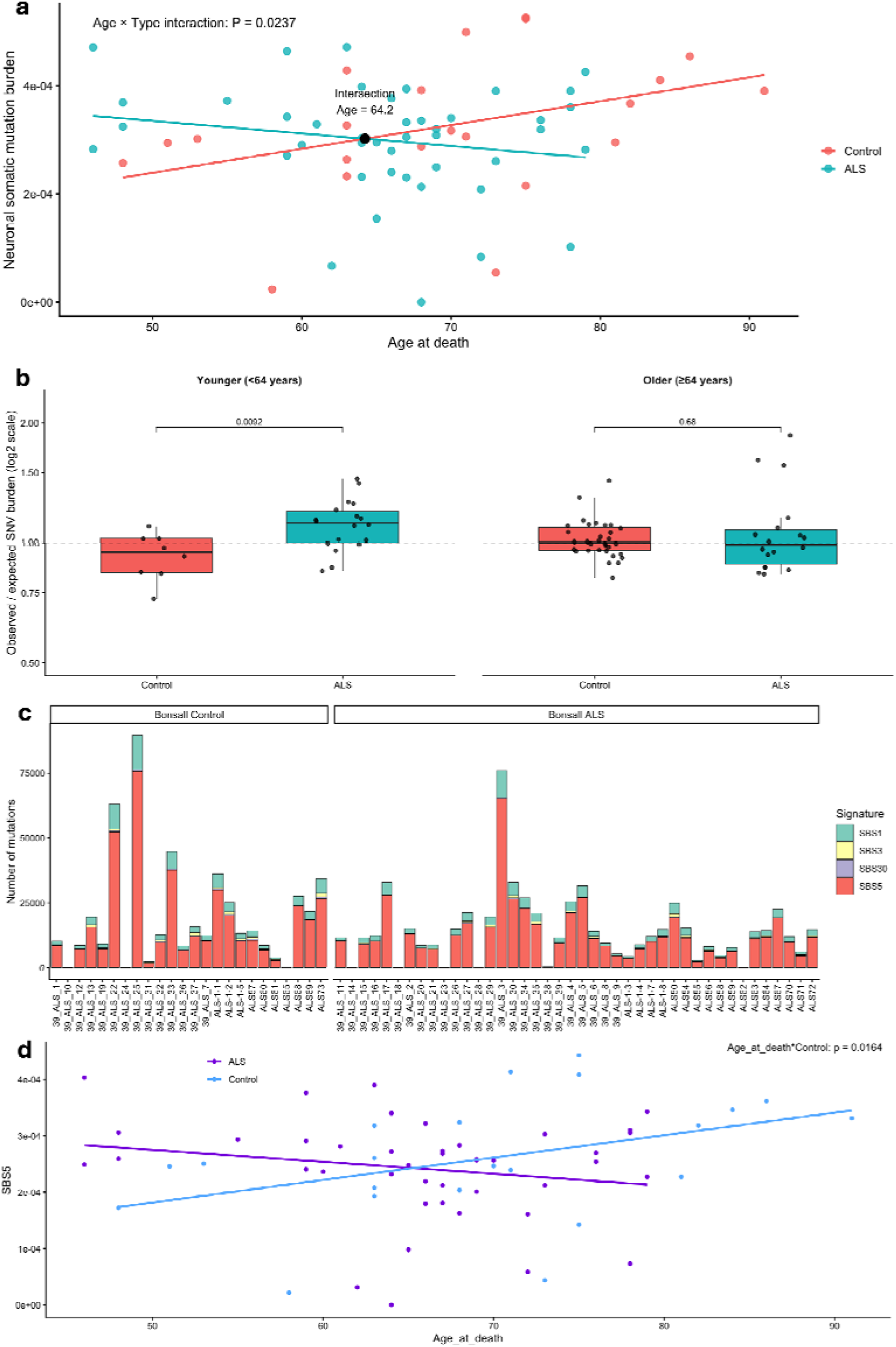
ALS alters the age-associated trajectory of neuronal somatic mutation burden. **a**, Association between age at death and neuronal somatic mutation burden in ALS and control samples from the Bonsall snATAC-seq dataset. Points represent individual samples, and lines indicate group-specific linear regression fits. The Age × disease-status interaction was significant (P = 0.024), indicating that the age-associated trajectories differed between ALS and control samples. The fitted regression lines intersected at an estimated age of 64.2 years. **b,** Age-stratified analysis of neuronal somatic SNV burden in the ALS single-cell PTA dataset. Left, age-corrected neuronal somatic SNV burden in ALS and control donors younger than 64 years. ALS neurons showed significantly higher somatic SNV burden than control neurons (Wilcoxon rank-sum test, P = 0.0092). Right, age-corrected neuronal somatic SNV burden in ALS and control donors aged 64 years or older, with no significant difference observed between groups (Wilcoxon rank-sum test, P = 0.68). **c,** COSMIC mutational signature exposures in neuronal sSNVs from ALS and control samples in the Bonsall snATAC-seq dataset. Stacked bars show the estimated number of mutations attributed to SBS1, SBS3, SBS30, and SBS5 in each sample. SBS5 accounted for the largest proportion of the fitted mutational burden across most samples. **d,** Association between age at death and neuronal SBS5 exposure in ALS and control samples. Points represent individual samples, and lines indicate group-specific linear regression fits. SBS5 increased with age in controls but decreased with age in ALS, resulting in a significant Age × disease-status interaction (P = 0.0164).

We hypothesized that preferential loss of neurons carrying a high somatic mutation burden during ALS progression could provide one explanation for the reversal of neuronal mutational burden observed between ALS and control samples at older ages. To test whether the burden of detectable neuropathology varied with donor age at death, we used neuronal expression of the STMN2 cryptic exon (CE) [55–57] as a proxy for TDP-43 dysfunction. The absolute number of STMN2 CE–positive neurons varied markedly across samples (Extended Data Fig. 4a), potentially reflecting biological heterogeneity and /or variable sensitivity for STMN2-CE detection. Nonetheless, the sample-level proportion of STMN2 CE–positive neurons were not associated with age at death in ALS samples (R = −0.042, P = 0.84; Extended Data Fig. 4b). While these findings do not fully exclude selective loss of highly mutated neurons in older patients, they do indicate that burden of pathology, as measured using STMN2 CE–positive neurons, does not appear to explain the age-dependent shift in neuronal somatic mutation burden.

Finally, to determine which mutational processes contributed to the ALS-associated trajectory shift, we decomposed neuronal somatic mutations into COSMIC mutational signatures using a constrained refitting strategy [31][40]. Four signatures: SBS1, SBS3, SBS5 and SBS30 were retained for downstream analysis after forward selection as described in methods (Fig. 2c, Method, Extended Data Fig. 2c). Among these signatures, SBS5 was the only component showing a nominally significant diagnosis-by-age interaction (P = 0.016; Figure 2c), whereas SBS1 showed borderline nominal evidence and SBS3/SBS30 were not significant (SBS1, P = 0.052, Extended data 5a-c). SBS5 is a clock-like signature that correlates with age across tissues [48], although its precise aetiology remains unresolved and may capture contributions from multiple endogenous [18,47] or environmental mutagenic processes [16–18]. Therefore, the SBS5 interaction suggests an association between ALS and a reshaping of a normal aging-associated mutational process.

High-resolution single-cell DNA sequencing has transformed the study of neuronal somatic mutagenesis, but disease-associated patterns remain difficult to generalize because existing studies are often constrained by limited donor numbers and numbers of profiled neurons [6,23,24,58]. Here, we used established 10X single-nucleus motor cortex cohorts to investigate neuronal somatic mutagenesis across a large number of ALS and control donors. Although sparse coverage restricts mutation detection to a limited genomic window, prior reports indicate that neuronal somatic mutations are enriched in regulatory and transcriptionally active regions [8,10,30,31]. Within this window, snATAC-derived variants showed several established features of neuronal somatic mutagenesis, including higher detectable burden in neurons, age-associated accumulation and recovery of the clock-like SBS5 signature. Thus, snATAC-seq does not provide a comprehensive measure of genome-wide mutation burden, but can provide an informative window for investigating cohort-level somatic mutational patterns.

Our principal finding was that ALS associated with an altered age-associated trajectory of neuronal somatic mutation burden rather than a uniform increase relative to controls. The fitted trajectories indicated higher burden in younger ALS donors, with the ALS-control difference diminishing and eventually reversing with age, not explained by loss of neurons with a higher burden of mutations. This pattern was predominantly associated with SBS5, which represented the largest fitted mutational component and showed the clearest diagnosis-by-age interaction. SBS5 is a clock-like signature that accumulates with age across human tissues, although its underlying aetiology remains unresolved and may encompass multiple endogenous and environmental processes [17,47,48]. Importantly, re-analysis of independent single-cell whole-genome sequencing data generated using PTA [26] recovered a similar age-dependent pattern, with ALS neurons exhibiting higher somatic SNV burden in younger donors, but not in older donors. Although this comparison was based on only six donors, concordance across distinct sequencing approaches supports the biological plausibility of the changing ALS-control relationship across age.

Several non-mutually exclusive processes could explain this pattern. ALS could alter the accumulation of SBS5-associated mutations during neuronal aging, or the preferential vulnerability of neurons carrying higher mutational burdens could instead change the population of surviving neurons sampled at older ages. We found no association between age at death and STMN2 cryptic-exon-positive neuronal burden, suggesting that the changing trajectory was not simply explained by greater TDP-43 pathology in older donors. However, selective loss of highly mutated neurons cannot be excluded. Patient heterogeneity could also contribute to this pattern. ALS is clinically and genetically heterogeneous [1], with major genetic subtypes differing in age at onset and disease trajectory [59]. Consequently, the genetic and clinical composition of ALS donors may differ across age groups.

A key question for future work is why the contribution of SBS5 differs across the ALS aging trajectory. Larger single-cell whole-genome studies will be required to validate this observation, determine whether it persists within different patient subgroups, and establish whether it extends beyond the restricted genomic regions sampled by snATAC-seq. However, larger post-mortem studies alone will remain cross-sectional and cannot determine whether ALS alters the accumulation of SBS5-associated mutations over time. Longitudinal human neuronal models therefore offer an important complementary approach. SBS5 is already known to be active in cultured human iPSCs, where it accounts for a substantial proportion of acquired somatic mutations [60], suggesting that these systems could provide an opportunity to study SBS5-associated mutation accumulation over time. However, whether iPSC-derived neurons reproduce the rate and spectrum of SBS5-associated mutation accumulation observed during neuronal aging in vivo remains unclear. Reprogramming through pluripotency also resets multiple features of cellular aging [61,62], potentially limiting the ability of iPSC-derived neurons to model donor-age-associated processes. Directly converted neurons retain more donor-age-associated characteristics [61], including aging hallmarks in directly reprogrammed human motor neurons [63], and may therefore provide a complementary approach. Long-term brain organoid models provide another emerging opportunity, with recent work showing that cortical organoids maintained for several years can reproduce aspects of the temporal maturation and aging of the human brain [64]. Additionally, given that patient-age-associated mutational patterns can be retained in patient-derived organoids, organoids derived from donors across different age groups may provide a useful model for investigating age-associated clock-like mutagenesis, including the accumulation of SBS5-associated mutations [65]. Together, these systems provide increasingly tractable opportunities to investigate the temporal dynamics of SBS5-associated somatic mutagenesis, while also highlighting the importance of establishing how closely their mutational processes reflect those operating in aging human neurons.

An important limitation of this study arises from the inherent characteristics of snATAC-seq, which provides sparse and nonuniform genomic coverage restricted primarily to accessible chromatin regions. Consequently, differences in neuronal recovery, sequencing depth, and callable genomic territory may influence somatic mutation detection and burden estimation. Furthermore, because chromatin accessibility and cellular composition may themselves be affected by disease-related biological processes, technical and biological contributions to the observed variation cannot be fully disentangled using the current data.

In conclusion, cohort-scale snATAC-seq revealed an age-dependent divergence in neuronal somatic mutation burden between ALS and controls, with SBS5 representing the major contributing mutational process. The concordant pattern observed in independent single-cell whole-genome data supports the biological plausibility of this finding, while its underlying mechanism remains unresolved. These results identify age as an important dimension of neuronal somatic mutagenesis in ALS and motivate larger single-cell genomic studies and longitudinal neuronal models to determine the processes underlying the SBS5-associated trajectory.

## Methods

### Data preprocessing

Raw FASTQ files were obtained from NCBI Li snATAC&snRNA: NCBI GEO under accession GSE219281 [38]; Bonsall snATAC&snRNA [39] ;Pineda snRNA:NCBI GEO under accession GSE174332 [40]. Bonsall snATAC&snRNA was provided by the authors [39]. snRNA-seq reads were processed and aligned using Cell Ranger v7.1.0 with default parameters and the 10X Genomics GRCh38 gene-expression reference, refdata-gex-GRCh38-2020-A. snATAC-seq reads were processed using Cell Ranger ATAC v2.1.0 with default parameters and the 10X Genomics GRCh38 ATAC/ARC reference, refdata-cellranger-arc-GRCh38-2020-A-2.0.0.

### Somatic variant calling using SComatic

Somatic variant calling was performed using SComatic, following the workflow recommended by the authors. Because our aim was to detect rare, cell-specific somatic mutations that accumulate with aging in non-neoplastic brain tissue, we modified several default parameters to increase sensitivity for low-frequency events. In step 2, which collects base-count information, --min_cc was reduced from 2 to 1. In step 4.2, which detects somatic mutations, --min_cells, --min_ac_cells and --min_cell_types were each set to 1. These settings were chosen to allow detection of variants supported by a single cell, consistent with the expected sparsity of age-associated somatic mutations in post-mitotic neuronal populations. Callable sites were identified using SComatic. For each dataset, the corresponding callable genomic regions were defined as the 10X-detectable genomic window. Unless otherwise stated, downstream analyses were performed within the 10X-detectable window of the snATAC-seq datasets. To minimize potential germline contamination while retaining candidate somatic variants in non-cancerous tissue under conditions of limited read depth, we applied a VAF threshold of <0.5. We selected this relatively permissive cutoff, rather than the more stringent VAF <0.25[66] threshold commonly used for somatic variant filtering, to maximize the retention of potential somatic mutations while excluding variants with VAFs more consistent with germline variation.

### Somatic variant simulation and benchmarking

To evaluate somatic variant calling performance in neurons, we used SomatoSim to simulate somatic variants in neuronal chromosome 21 regions. Simulations were performed using two representative samples from Li’s snATAC: one C9orf72 ALS sample, A2, and one control sample, C5. Simulated variants were generated across accessible neuronal regions and stratified by variant allele frequency according to previously defined categories [50]: early arising somatic SNVs, defined as VAF ≥ 0.05 and < 0.25; ultralow-frequency somatic SNVs, defined as VAF < 0.01; germline SNVs, defined as VAF > 0.25 and < 1.00; somatic SNVs, defined as VAF < 0.25; and combined germline and somatic SNVs, defined as VAF < 1.00. We simulated several VAF scenarios rather than mutually exclusive biological classes.

The simulated datasets were processed using the same SComatic workflow described above. Because true-negative somatic variants cannot be unambiguously defined in this synthetic setting, benchmarking focused on sensitivity, calculated as the proportion of simulated variants recovered by the variant calling pipeline. Sensitivity was evaluated across read depth and VAF strata to define the effective callable window of 10X multiome data.

### Statistical Analyses

Unless otherwise indicated, all statistical analyses were performed using R version 4.5.2.

Neuronal somatic mutation burden was modeled using linear regression with age at death, diagnosis group (case vs control) and their interaction as predictors:

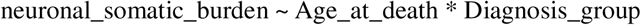

The interaction term tested whether the age-associated trajectory of neuronal somatic mutation burden differed between ALS and controls. A significant interaction was interpreted as evidence that ALS modifies the relationship between age and neuronal somatic mutation burden.

### Mutational signature analysis

Mutational signature analysis was performed using the R package MutationalPatterns v3.20.1. To mitigate against overfitting, we constructed a curated panel of 20 COSMIC mutational signatures previously reported in brain or brain-related somatic mutation studies [12,24,40–45]. Signature refitting was performed using a constrained non-negative least-squares framework. Candidate signatures were refined using forward stepwise selection, following strategies used in previous mutational signature studies [31][40].

This procedure retained four signatures for snATAC Bonsall and Li all-cell-type analyses and snATAC Bonsall neuronal analyses. Seven signatures were retained for snRNA Bonsall, Li and Pineda analyses. For each neuronal sample, mutational profiles were decomposed into the relative contributions of 20 preselected COSMIC mutational signatures, expressed as percentages, using signature refitting mentioned above. The relative contribution of each signature was then multiplied by the total neuronal somatic mutation burden to derive its signature-specific burden. Signature-specific burden estimates were subsequently used to assess the contribution of individual mutational processes to the age-by-diagnosis interaction.

### STMN2 cryptic exon-defined diseased neurons

We used the STMN2 cryptic exon (CE)-positive neuron list originally reported by Bonsall [39] and provided in [67] to identify diseased neurons for downstream analyses.

## Data availability

Full metadata, including mutational burden of different cell types, individual’s phenotypes, individuals age at death.etc from this study are available under the following link:

https://github.com/Li628/ALS_multiomics_somatic_mutation

## Code availability

Codes used for analyses and plots in this study are available under the following link: https://github.com/Li628/ALS_multiomics_somatic_mutation

## Acknowledgements

This work was sponsored by NWO-Domain Science for the use of supercomputer facilities.

## Extended Data

**Extended Data Fig. 1.**
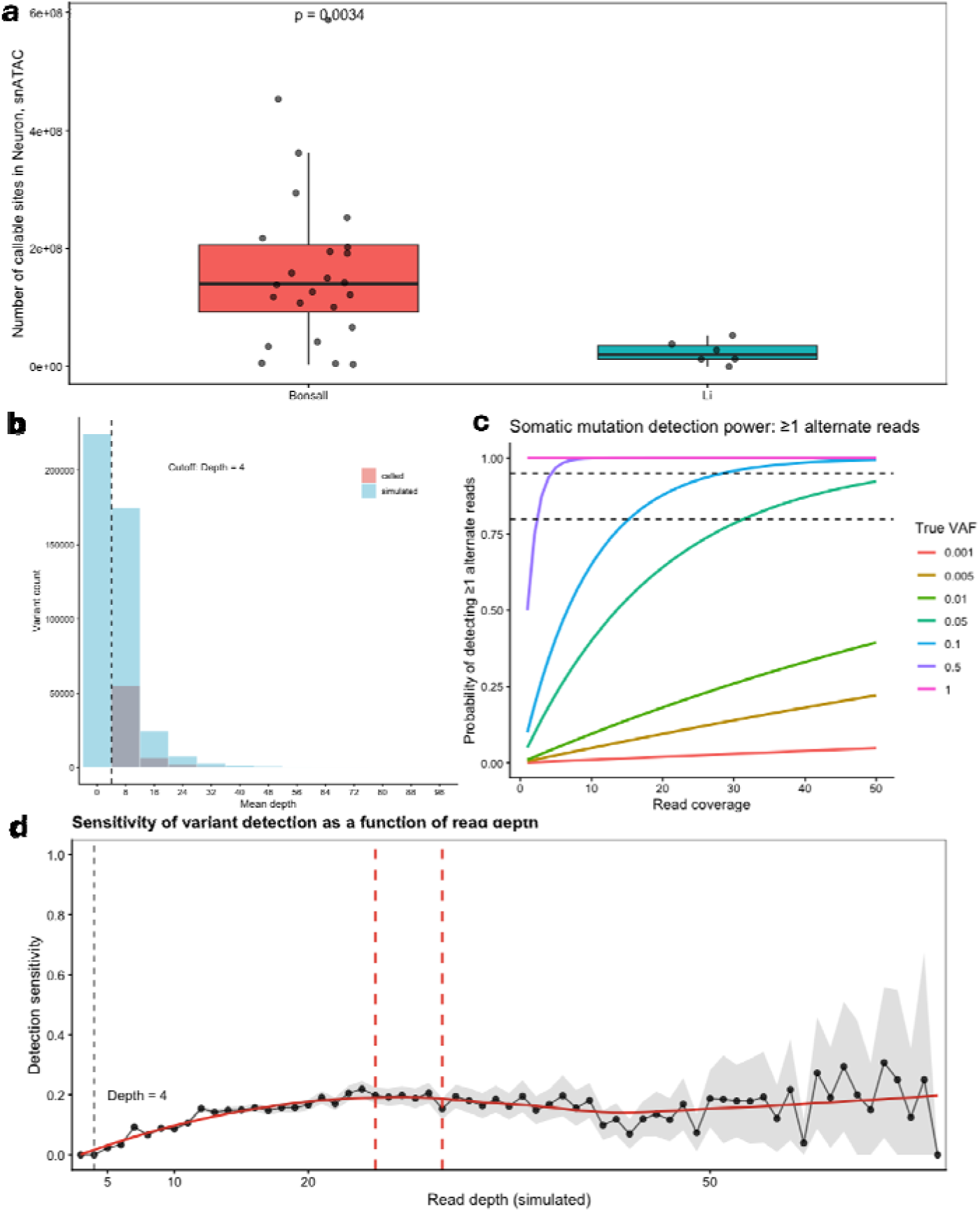
Neuronal callable-site coverage differs between the Bonsall and Li snATAC-seq datasets. **a**, Distribution of the number of callable sites in neurons across samples from the Bonsall and Li snATAC-seq datasets. Bonsall samples had significantly more neuronal callable sites than Li samples (P=0.0034, two-sided Wilcoxon rank-sum test). Points represent individual samples; boxes show the median and interquartile range, and whiskers extend to 1.5 times the interquartile range. **b,** Distribution of read depth for simulated variants and variants recovered by the calling pipeline. The vertical dashed line indicates the minimum read-depth threshold of four used to define mutation-detectable sites. c, Theoretical somatic mutation detection probability as a function of read depth and variant allele frequency (VAF). The probability of observing at least one alternate read was calculated across read depths of 1–50 using a binomial model for variants with true VAFs ranging from 0.001 to 1. Detection probability increased with read depth but was strongly dependent on VAF, with low-VAF variants remaining difficult to detect even at relatively high coverage. Dashed horizontal lines indicate 80% and 95% detection probabilities. d, Sensitivity of simulated-variant recovery as a function of read depth. Points indicate the observed detection sensitivity at each simulated depth, the black line connects observed values, the red line shows the smoothed trend and the grey shaded area indicates the 95% confidence interval. The grey dashed line marks the minimum read-depth threshold of five.

**Extended Data Fig. 2.**
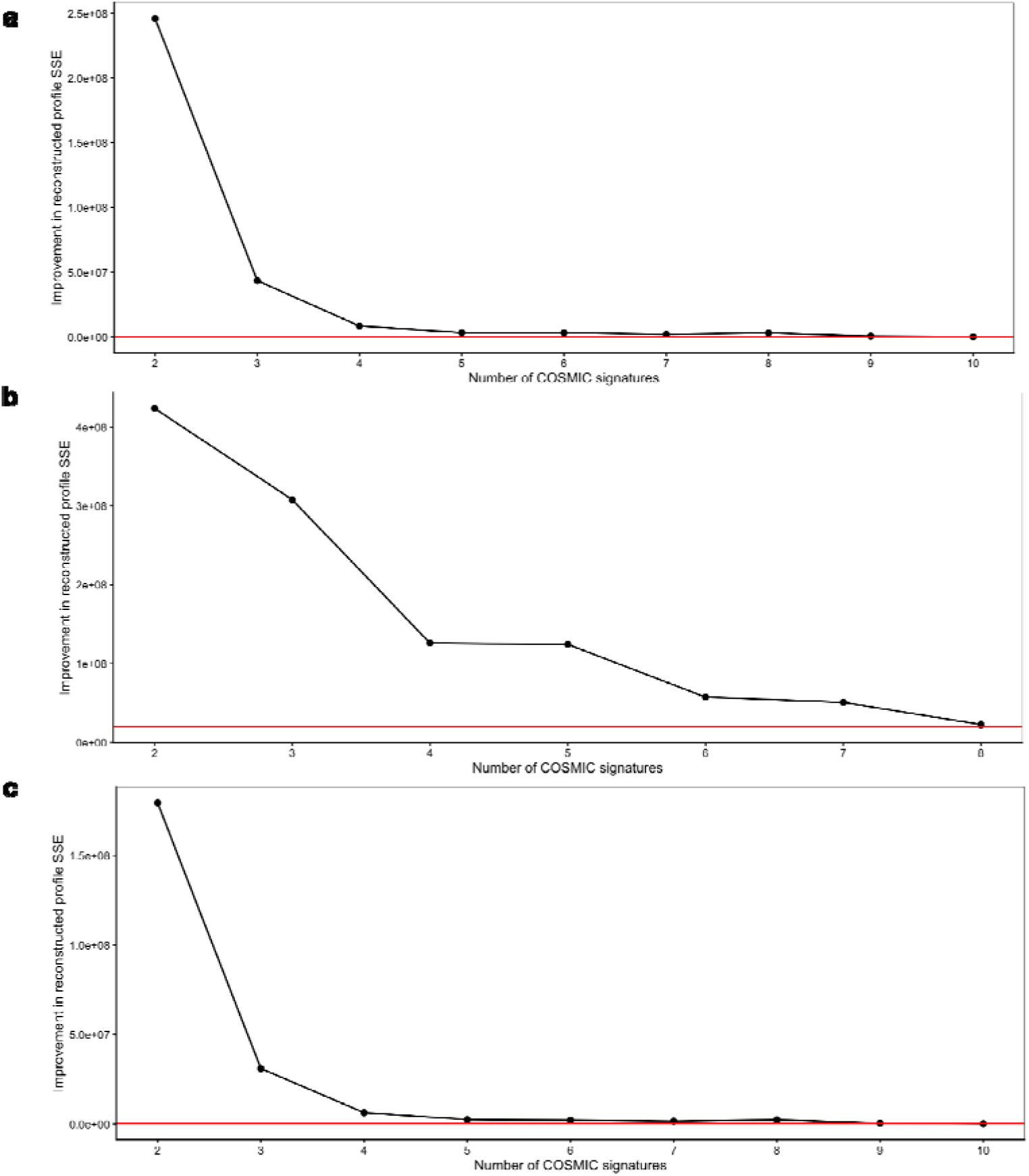
Dataset-specific selection of COSMIC mutational signatures using stepwise reconstruction-error improvement. **a**, Stepwise signature selection using all sSNVs detected across the snATAC-seq datasets. The red horizontal line indicates the empirical stopping threshold of 200,000 in SSE improvement. Signature addition was stopped once the improvement fell below this threshold, resulting in the retention of four signatures. **b,** Stepwise signature selection using all sSNVs detected across the snRNA-seq datasets. The red horizontal line indicates the empirical stopping threshold of 20,000,000 in SSE improvement. Six signatures were retained for subsequent refitting. **c,** Stepwise signature selection using neuron-specific sSNVs detected across the snATAC-seq datasets. The red horizontal line indicates the empirical stopping threshold of 200,000 in SSE improvement. Four signatures were retained for downstream analyses.

**Extended Data Fig. 3.**
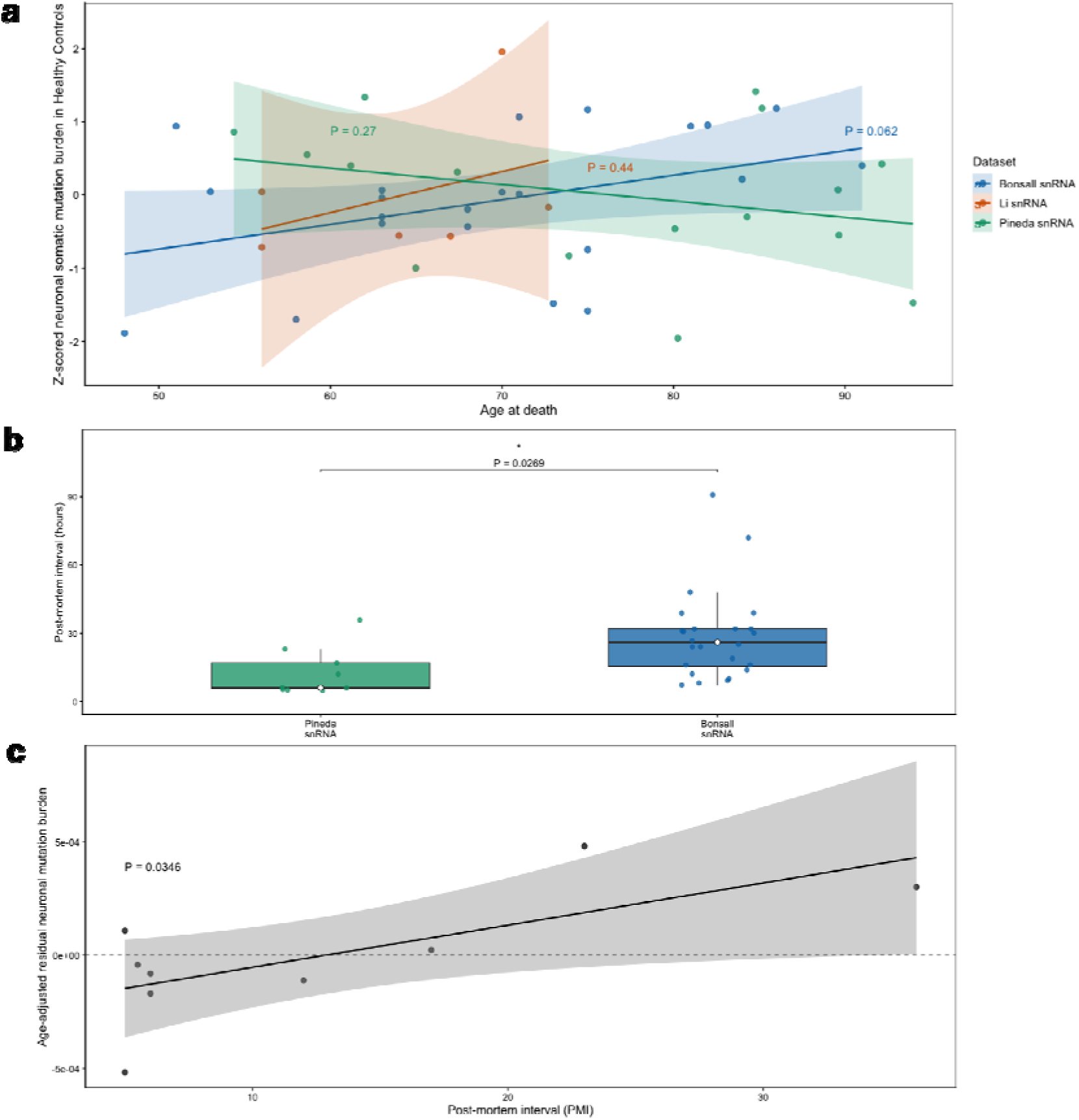
Neuronal somatic mutation burden analyses in snRNA-seq datasets and assessment of post-mortem interval as a technical covariate. **a**, Association between age at death and z-scored neuronal somatic mutation burden in three snRNA-seq datasets: Bonsall (P=0.062), Li (P=0.44), and Pineda (P=0.27). The Bonsall and Li datasets showed positive age-related trends, whereas the Pineda dataset showed a negative trend. **b,** Distribution of PMI values among healthy control samples from the Pineda and Bonsall snRNA-seq datasets. PMI differed significantly between the two datasets (P=0.0269). PMI information was unavailable for healthy control samples in the Li dataset. **c,** Association between post-mortem interval (PMI) and age-adjusted neuronal somatic mutation burden in healthy control samples from the Pineda snRNA-seq dataset. PMI was significantly associated with the age-adjusted residual burden (P=0.0346), supporting its inclusion as a technical covariate.

**Extended Data Fig. 4.**
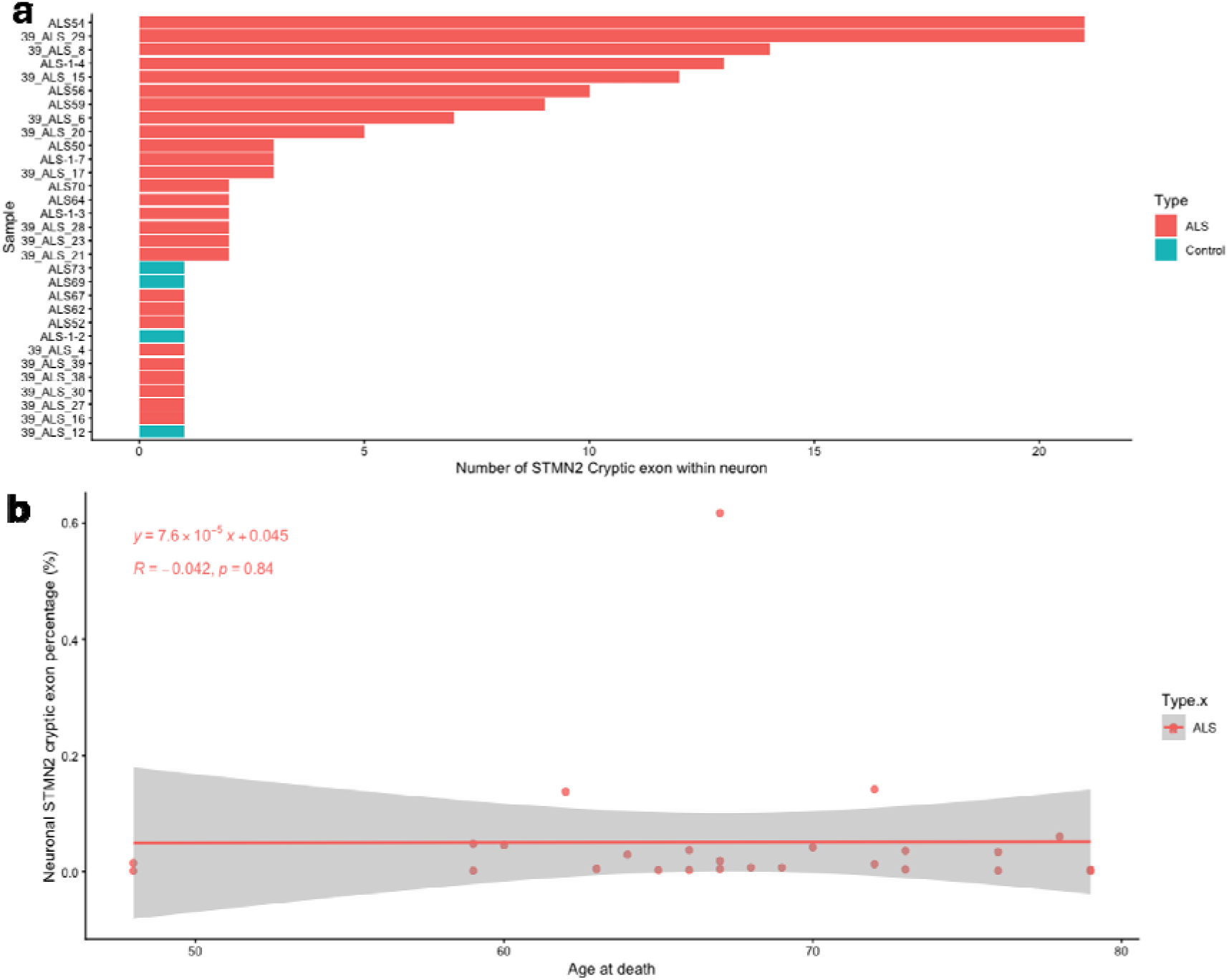
STMN2 cryptic exon–defined neuronal involvement is not associated with age at death in the Bonsall snATAC-seq dataset. **a**, Number of STMN2 cryptic exon–positive neurons detected per sample in the Bonsall snATAC-seq dataset. Bars are colored by disease status. **b,** Association between age at death and the proportion of neurons containing the STMN2 cryptic exon in ALS samples from the Bonsall snATAC-seq dataset. No significant association was observed (R=−0.042, P=0.84).

**Extended Data Fig. 5.**
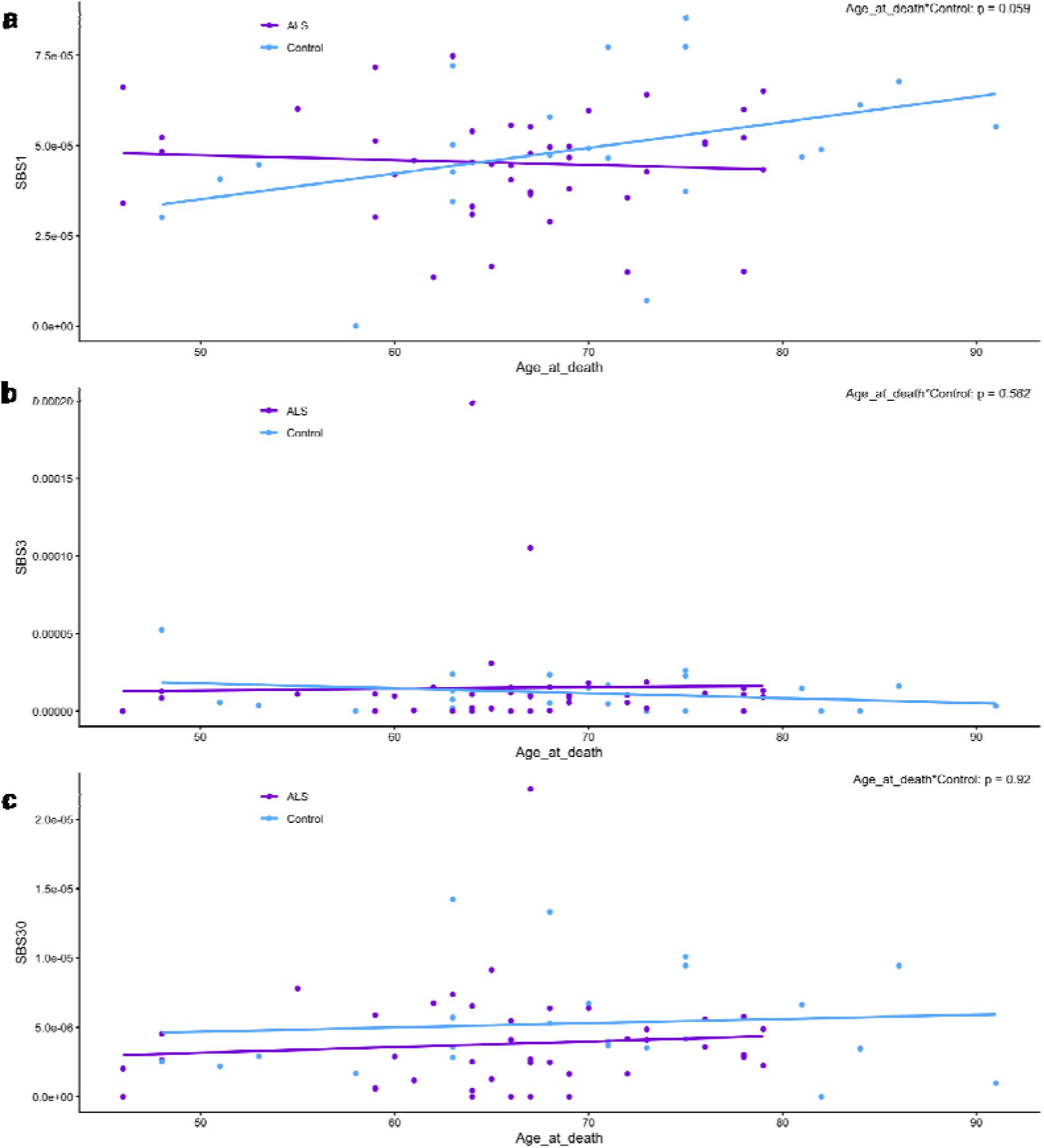
Age-related trajectories of additional COSMIC signature exposures in neuronal sSNVs from the Bonsall snATAC-seq dataset. **a**, Association between age at death and neuronal SBS1 exposure in ALS and control samples. The Age × disease-status interaction was borderline significant (P=0.059). **b,** Association between age at death and neuronal SBS3 exposure in ALS and control samples. No significant Age × disease-status interaction was observed (P=0.582). **c,** Association between age at death and neuronal SBS30 exposure in ALS and control samples. No significant Age × disease-status interaction was observed (P=0.920).

